# Golgi retention of KIT in gastrointestinal stromal tumour cells is phospholipase D activity-dependent

**DOI:** 10.1101/2025.03.02.640696

**Authors:** Yuuki Obata, Miyuki Natsume, Isamu Shiina, Tsuyoshi Takahashi, Toshirou Nishida

## Abstract

A constitutively active mutant of the receptor protein-tyrosine kinase KIT is a major cause of gastrointestinal stromal tumours (GISTs). Recently, we discovered that during biosynthetic transport, the KIT mutant (KIT^mut^) is retained in the Golgi/*trans-*Golgi network (TGN), where it activates downstream molecules. This retention is dependent on the phospholipase Cγ2–protein kinase D2–PI4 kinase IIIβ (PLCγ2–PKD2–PI4KIIIβ) pathway, which KIT^mut^ activates at the Golgi/TGN. The activated cascade aberrantly recruits GGA1 and the γ-adaptin subunit of AP1, resulting in KIT^mut^ retention in the Golgi/TGN. However, the precise mechanisms, including the mediators and effectors in the pathway, remain unclear. In humans, the phosphatidic acid-generating enzymes, phospholipase D1 (PLD1) and PLD2, are known downstream proteins of PKD. In the presence of the PLD inhibitor CAY10594, KIT^mut^ is released from the Golgi/TGN and subsequently degraded in lysosomes, leading to signal inactivation. Knockdown experiments indicated that PLD2 plays a role in KIT^mut^ retention. KIT^mut^ activates PLD2 through PKD2, but not PI4KIIIβ, for Golgi/TGN retention. PLD activity is required for the association of γ-adaptin with GGA1. Therefore, the KIT–PLCγ2–PKD2 pathway separately activates PLD2 and PI4KIIIβ to recruit γ-adaptin and GGA1. Collectively, these results suggest that KIT^mut^ retention is dependent on the activation of the PLCγ2–PKD2–PLD2 cascade in GIST.

## Introduction

The stem cell factor (SCF) receptor, KIT, is a type III receptor protein-tyrosine kinase (RTK) that is primarily located in the plasma membrane (PM) of the interstitial cells of Cajal in the gastrointestinal tract, as well as in haematopoietic cells, mast cells, melanocytes, and germ cells in the testis^1–3^. Upon SCF binding to the PM, KIT dimerises, resulting in enhanced protein tyrosine kinase activity. This, in turn, activates several signalling molecules, such as AKT, extracellular signal-regulated kinase, phospholipase Cγ (PLCγ), and signal transducer and activator of transcription (STAT) proteins, leading to cell growth, survival, migration, and differentiation^1^. Consequently, the constitutive activation of KIT through genetic abnormalities, such as point mutations and in-frame deletions, plays a critical role in cancerogenesis. Indeed, most cases of mast cell leukaemia in adult humans, as well as patients with gastrointestinal stromal tumours (GISTs), harbour KIT mutations^4–7^. As a result, imatinib, a KIT inhibitor (tyrosine kinase inhibitor (TKI)), has been used to treat patients with advanced GISTs. However, 2–3 years after the administration of imatinib, TKIs become ineffective in the treatment of advanced GISTs^8–10^. In most cases, mutations conferring imatinib resistance are found in the *KIT* gene.

Previously, we reported that, unlike normal KIT, mutant KIT (KIT^mut^) in leukaemia cells and GIST is localised to intracellular compartments where it can be autophosphorylated and activated^11–15^. Moreover, the localisation of the mutant in leukaemia cells is notably different from that in GIST cells; in leukaemia cells, KIT^mut^ accumulates in the endosome–lysosome compartments^11, 13, 15^, whereas in GIST cells, it is retained in the Golgi/*trans*-Golgi network (Golgi/TGN) area during biosynthesis^12, 14^. In both cases, KIT^mut^ localises to intracellular compartments depending on its tyrosine kinase activity^11, 12, 15^. We recently demonstrated that KIT^mut^ retention depends on protein kinase D2 (PKD2) activity in GIST cells^16^. The mediator between KIT^mut^ and PKD2 is PLCγ2. It was suggested that the KIT–PLCγ2–PKD2 pathway activates the phosphatidylinositol 4-phosphate-producing (PI4P-producing) enzyme PI4 kinase IIIβ (PI4KIIIβ), leading to the aberrant recruitment of Golgi-localised, γ-ear-containing, ARF-binding family of protein 1 (GGA1) and an adaptor protein-1 complex (AP1) component γ-adaptin. Subsequently, KIT itself is retained in the Golgi/TGN area. These findings raised several questions: How do the aberrantly recruited AP1 and GGA1 trap KIT in the Golgi/TGN?; Are there PKD2 effectors other than PI4KIIIβ? These questions remain unresolved.

Several studies have suggested that PKD enhances phosphatidylcholine-selective phospholipase D (PC-PLD; hereafter referred to as PLD) activity to generate phosphatidic acid (PA) from PC on the cytoplasmic face of the PM and Golgi/TGN membrane^17–19^. Under normal conditions, PA plays a key role in forming transport carriers in the Golgi/TGN membrane^20–25^. It creates a negative curvature at the budding site of membrane carriers due to its cone-like shape and provides a hydrophobic zone as a scaffold region for functional proteins^26–28^. Human PLDs consist of six structurally related proteins: PLD1, PLD2, PLD3, PLD4, PLD5, and PLD6. PLD1 and PLD2 are crucial for membrane carrier formation through PLD activity, while PLD3, PLD4, and PLD5 lack canonical phospholipase activity and may possess endonuclease activity in lysosomes and the endoplasmic reticulum (ER)^28, 29^. PLD6 is localised in mitochondria to produce PA from cardiolipins. Therefore, we hypothesised that the KIT–PLCγ2–PKD2 pathway activates PLD1 and/or PLD2, and that aberrant PA production leads to the mutant receptor’s retention in the Golgi/TGN in GIST cells.

The present study aimed to determine the requirement of PLD activity for Golgi/TGN retention of KIT^mut^ in GIST cells by investigating the relationship between the PLCγ2–PKD2 pathway and PLDs using PLD inhibitors, knockdown experiments, immunofluorescence confocal microscopic analysis, and biochemical assays.

## Results

### KIT^mut^ is retained in the Golgi/TGN region depending on PLD activity in GIST cells

Previous studies have shown that PLD and PI4KIIIβ are activated downstream of PKD^17–19^. We hypothesised that PLD activity was also involved in the retention of KIT^mut^ in the Golgi/TGN region of GIST cells. To evaluate this hypothesis, we treated GIST-T1 cells, which endogenously express a constitutively active KIT^Δ560–578^ mutant (imatinib-sensitive)^30^, with CAY10594 (an inhibitor of PLD activity)^31, 32^ and examined whether PLD inhibition mimicked the inhibition of PKD2 on KIT localisation and growth signalling. First, we performed an immunofluorescence confocal microscopic assay with anti-KIT and anti-golgin97 (TGN marker) antibodies in GIST-T1 cells treated with 20 µM CAY10594. As shown in Figure 1a, KIT was found in the Golgi/TGN area together with golgin97 under normal conditions (upper panels), whereas it was absent in CAY10594-treated cells (lower panels). This effect of CAY10594 was similar to that of the PKD inhibitor CRT0066101 in releasing KIT from the Golgi/TGN region (compare Fig. 1a with Suppl. Fig. 1a). Immunoblotting revealed that CAY10594 treatment decreased KIT protein levels in a time- and dose-dependent manner (Fig. 1b, upper panels). This effect was similar to that of PKD2 inhibition by CRT0066101 and small interfering RNA (siRNA), which induce the degradation of KIT in lysosomes^16^. Under PLD-inhibited conditions, the KIT downstream proteins, AKT and STAT5, were dephosphorylated and inactivated (Fig. 1b). Treatment with other PLD inhibitors, such as 1-butanol^20, 33, 34^, gave similar results (Fig. 1c). These results suggest that PLD activity is required for KIT^mut^ signalling by trapping the mutant in the Golgi/TGN region. Importantly, neither CAY10594 nor 1-butanol decreased phospho-PKD2 S876 (pPKD2^S876^) levels (Fig. 1b, c), indicating that PKD2 is not a downstream molecule of PLD2. Next, we examined the effect of PLD inhibition with CAY10594 in imatinib-resistant GIST cell lines, GIST-R9 (*KIT^Δ560–578,D820V^*) and GIST430 cells (*KIT^Δ560–576,V654A^*). Although higher concentrations of the compound were required, GIST-R9 and GIST430 cells showed results similar to those of CAY10594-treated GIST-T1 cells; the protein levels of KIT decreased, AKT and STAT5 were dephosphorylated, and pPKD2^S876^ was not reduced (Fig. 1d; Suppl. Fig. 1b). Taken together, these results suggest that PLD, a presumed PKD effector, plays a key role in Golgi/TGN retention of KIT^mut^.

**Figure 1.**
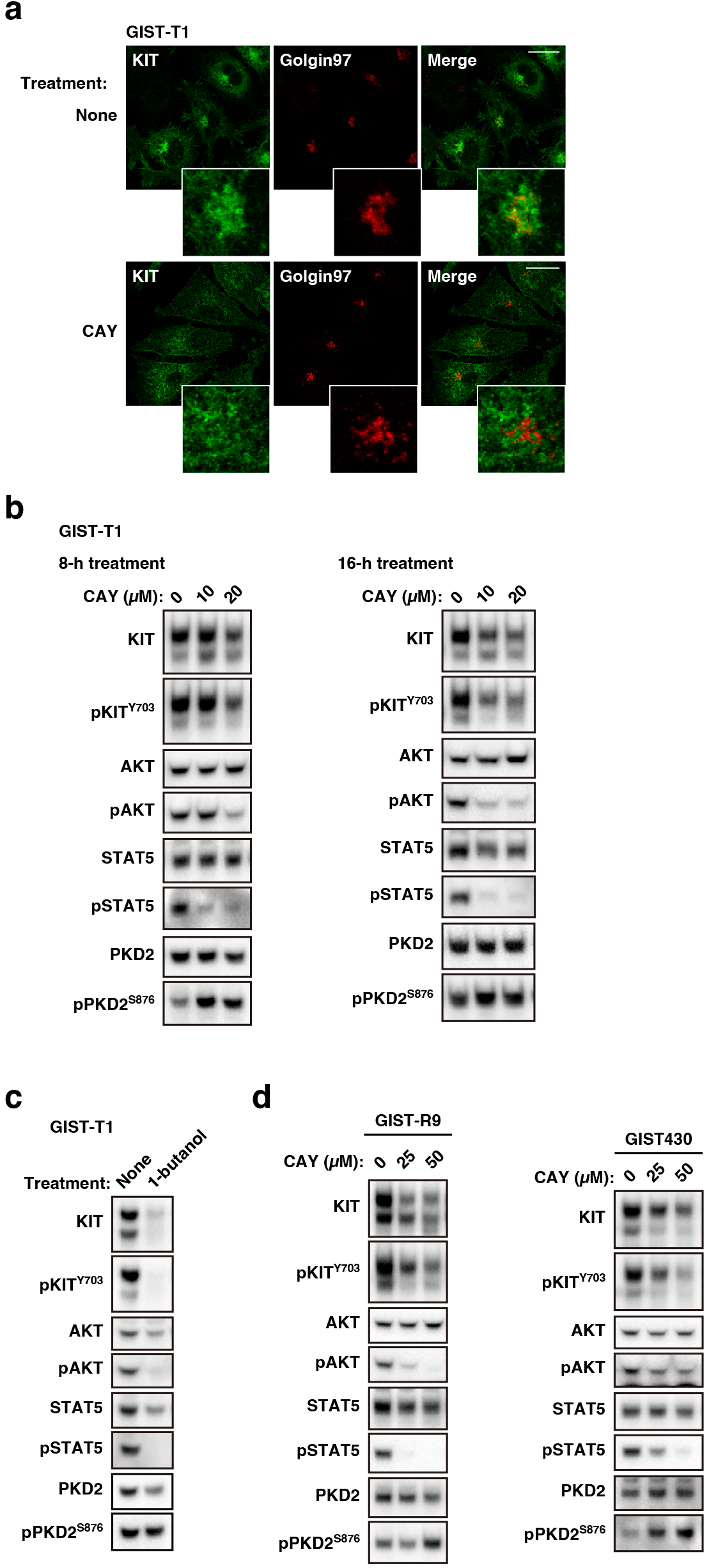
KIT^mut^ is trapped in the Golgi/TGN area in a manner dependent on PLD activity in GIST cells. **(a)** GIST-T1 cells were treated with 20 *µ*M CAY10594 (CAY, a PLD inhibitor) for 4 h, then immunostained for KIT (green) and golgin97 (Golgi marker, red). Insets show magnified images of the Golgi/TGN area. Scale bars, 20 *µ*m. Note that KIT disappeared from the Golgi/TGN area in the presence of CAY10594. **(b)** GIST-T1 cells were treated with CAY10594 for the indicated periods and then immunoblotted. pKIT^Y703^, phospho-KIT Y703; pPKD2^s876^, phospho-PKD2 S876. Note that the release of KITmut from the Golgi/TGN area by CAY10594 treatment inactivated AKT and STATS without decreasing pPKD2^s876^. **(c)** GIST-T1 cells were treated with 1% 1-butanol (inhibits PLD activity) for 8 h and then immunoblotted. **(d)** lmatinib-resistant cell lines, GIST-R9 cells (left) and GIST430 cells (right), were treated with CAY10594 for 8 h and then immunoblotted.

### KIT^mut^ released from the Golgi/TGN migrates to lysosomes in GIST cells

Recently, we reported that in PKD2-inhibited GIST cells, KIT^mut^ is released from the Golgi/TGN region into the PM and is subsequently degraded in lysosomes^16^. To examine whether CAY10594-induced KIT reduction was lysosome-dependent, we treated GIST-T1 cells with CAY10594 and ammonium chloride (NH_4_Cl), which inhibits lysosomal proteases^35^, followed by immunoblotting. Figure 2a shows that CAY10594 alone decreased the protein level of KIT, whereas the level of KIT was restored in cells treated with CAY10594 and NH_4_Cl. The levels of transferrin receptor, which circulates between the PM and endosomes^36^, were not affected by these treatments, suggesting that CAY10594 specifically acts on KIT^mut^ trafficking. In support of these results, immunofluorescence data showed that in the presence of CAY10594 and NH_4_Cl, KIT was found in the lysosome-associated membrane protein 1-positive area (Fig. 2b). CAY10594, therefore, has an effect similar to that of PKD2 inhibition on the distribution of KIT^mut^ in GIST cells. These results suggest that KIT^mut^ is degraded in lysosomes after CAY10594-induced release from the Golgi/TGN region.

**Figure 2.**
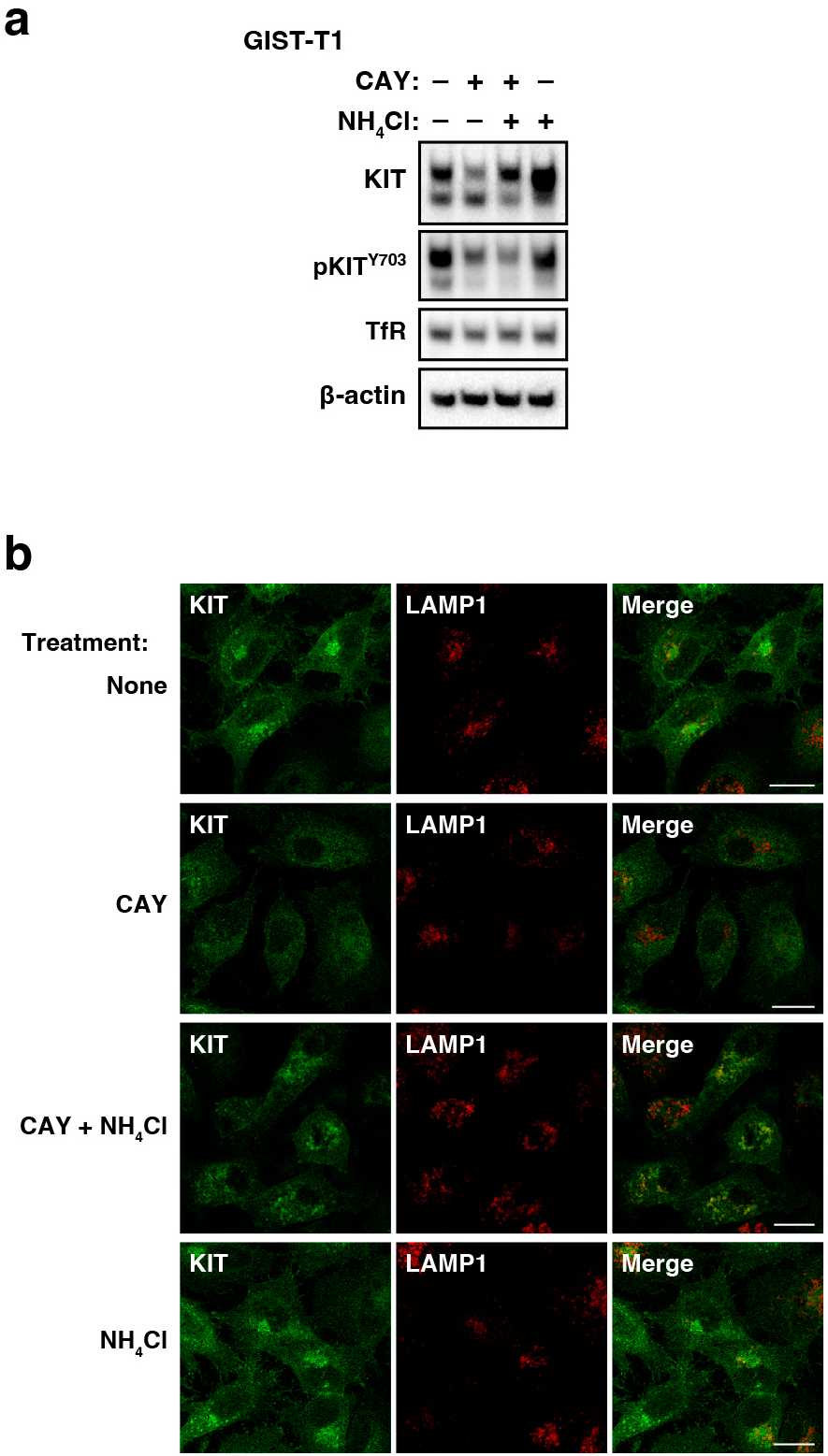
CAY10594-induced reduction of KIT is due to lysosomal degradation. **(a, b)** GIST-T1 cells were treated with 20 *µ*M CAY10594 (CAY, a PLD inhibitor) and/or 20 mM NH4CI (inhibits lysosomal proteases) for **(a)** 16 h or **(b)** 4 h. **(a)** Lysates were immunoblotted for KIT, phospho-KIT Y703 (pKIT^Y703^), transferrin receptor (TfR, a recycling endosome protein), and β-actin. **(b)** Cells were immunostained with KIT (green) and LAMP1 (lysosome-associated membrane protein 1, red). Scale bars, 20 *µm.* Note that KIT was found in lysosomes in GIST cells treated with CAY10594 plus NH_4_CI.

### PLD2 knockdown phenocopies the effect of CAY10594 treatment in GIST-T1 cells

Mammalian PLDs comprise six structurally related proteins: PLD1, PLD2, PLD3, PLD4, PLD5, and PLD6^29,37^. PLD1 and PLD2 have PA-producing PLD activity not only in the PM but also in the Golgi/TGN, whereas no canonical PLD activity of PLD3, PLD4, or PLD5 has been reported^29^. PLD6 hydrolyses cardiolipin to generate PA in the mitochondria and is called mito-PLD. Therefore, we examined whether the knockdown of PLD1 or PLD2 phenocopied the effect of chemical inhibition of PLD in GIST. In PLD2-knockdown GIST-T1 cells, KIT reduction, a sign of release from the Golgi/TGN region, and suppression of KIT signalling were observed (Fig. 3a). Similar to the CAY10594 treatment, PLD2 knockdown did not decrease the expression of pPKD2^S876^. PLD1 protein expression was not detected in this cell line; thus, PLD1 siRNA had no effect on KIT retention or growth signalling. Therefore, PLD2 plays a pivotal role in Golgi/TGN retention of KIT in GIST-T1 cells. At this stage, we could not determine whether PLD1 was important for KIT retention. Next, we investigated whether PLD2 is downstream of KIT^mut^. As shown in Figure 3b, imatinib, a KIT tyrosine kinase inhibitor, markedly decreased the phosphorylation of PLD2 both at residues Y169 and Y511, which are signs of PLD2 activation. Furthermore, PLD2 coimmunoprecipitated with KIT and vice versa, and this interaction was suppressed by imatinib treatment (Fig. 3c). These results suggest that PLD2 is located downstream of KIT^mut^ in GIST-T1 cells.

**Figure 3.**
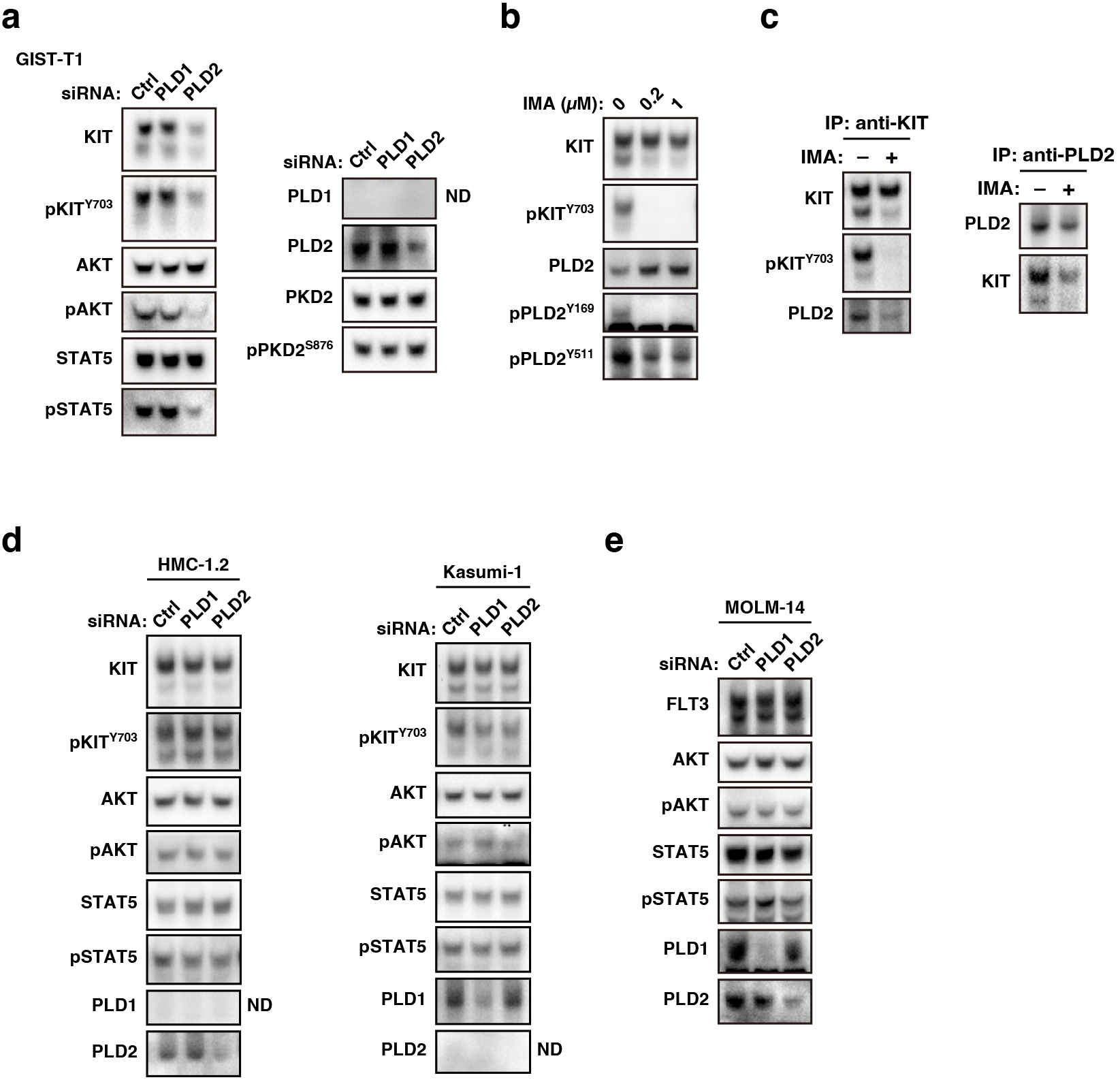
PLD2 is a downstream molecule of KIT^mut^ in GIST-T1 cells. **(a)** GIST-T1 cells were transfected with PLD1-targeted siRNA or PLD2-targeted siRNA and cultured for 30 h. Lysates were immunoblotted with the indicated antibodies. pKIT^Y703^, phospho-KIT Y703; pPKD2^s876^, phospho-PKD2 S876. ND, not detected. Note that PLD2 knockdown phenocopied CAY10594 treatment in GIST-T1 cells. **(b, c)** GIST-T1 cells were treated with imatinib (IMA, a KIT tyrosine kinase inhibitor) for 8 h. (b) Lysates were immunoblotted. pPLD2^Y169^, phospho-PLD2 Y169; pPLD2^Y511^, phospho-PLD2 Y511. **(c)** KIT (left) or PLD2 (right) in cells treated with 200 nM imatinib was immunoprecipitated. The immunoprecipitates were immunoblotted. IP, immunoprecipitation. **(d, e)** Kasumi-1 (AML, *KIT^N822K^),* HMC-1.2 (MCL, *KIT^D816v^),* or MOLM-14 cells (AML, *FLT3^ITD^)* were transfected with PLD1-targeted siRNA or PLD2-targeted siRNA and cultured for 48 h. Lysates were immunoblotted with the indicated antibodies. ND, not detected.

Next, we examined the subcellular localisation of PLD2. In our immunofluorescence assay, we were unable to detect endogenous PLD2, probably due to the low ability of the antibodies. Transiently transfected MYC-tagged PLD2 was found not only in the Golgi/TGN region but also in other endomembranes (Suppl. Fig. 2), indicating that a small fraction of PLD2 is associated with KIT^mut^ in GIST cells.

We next examined whether PLD knockdown had a similar effect on RTK^mut^ expression in leukaemia cells as on KIT^mut^ expression in GIST-T1 cells. We previously reported that KIT^mut^ in AML cells (Kasumi-1) and MCL cells (HMC-1.2) flows normally from the Golgi/TGN to the PM and immediately moves to the endosome–lysosome compartments^11, 13, 15^. We were unable to determine the expression of PLD2 and PLD1 in Kasumi-1 and HMC-1.2 cells, respectively. As shown in Figure 3d, compared with KIT in GIST-T1 cells, KIT in Kasumi-1 and HMC-1.2 was not decreased by PLD knockdown. In addition, we recently showed that in AML cells, the FLT3 internal tandem duplication (FLT3-ITD) mutant is retained in the Golgi/TGN, where it initiates the activation of AKT^38^. It activates STAT5 in the ER^38, 39^, suggesting that FLT3-ITD signalling occurs during its biosynthetic trafficking. The knockdown of PLD1 or PLD2 did not affect the protein levels of FLT3 or AKT activation in the AML cell line MOLM-14 (Fig. 3e). Furthermore, FLT3-ITD-dependent STAT5 activation in the ER was unaffected in PLD-knockdown cells. This supports our findings that Golgi/TGN retention in FLT3-ITD is a PKD2-independent manner^16^. Collectively, these results suggest that PLD activity is specifically involved in Golgi/TGN retention of KIT^mut^ in GIST cells.

### KIT–PLCγ2–PKD2 separately activates PI4KIIIβ and PLD2

Finally, we asked where PLD2 is located in the KIT–PLCγ2–PKD2–PI4KIIIβ–AP1–GGA1 pathway. Figure 4a shows that the knockdown of PLCγ2 as well as PKD2 reduced the pPLD2^Y511^, supporting previous reports that PLDs are downstream molecules of PKD^17–19^. On the other hand, pPLD2^Y511^ was maintained in PI4KIIIβ-knocked down cells compared with PKD2-knocked down cells, indicating that PLD2 is not a downstream molecule of PI4KIIIβ. Next, we examined PI4P levels in the Golgi/TGN region. Imatinib (KIT inhibitor), CRT0066101 (PKD inhibitor), and PIK-93 (PI4KIIIβ inhibitor) decreased PI4P in the Golgi/TGN region (Fig. 4b), consistent with our previous report. On the other hand, PI4P was found in the Golgi/TGN area in CAY10594 (PLD inhibitor). Our recent study showed that activation of the KIT–PLCγ2–PI4KIIIβ pathway is a cause of the association of γ-adaptin and GGA1. As shown in Figure 4c, inhibition of PI4KIIIβ, PKD2, and PLD2 decreased γ-adaptin interaction with GGA1. Collectively, these results suggest that PKD2 separately activates PLD2 and PI4KIIIβ to recruit AP1 and GGA1. Both PI4KIIIβ-produced PI4P and PLD2-produced PA probably contribute to the association of AP1 with GGA1.

**Figure 4.**
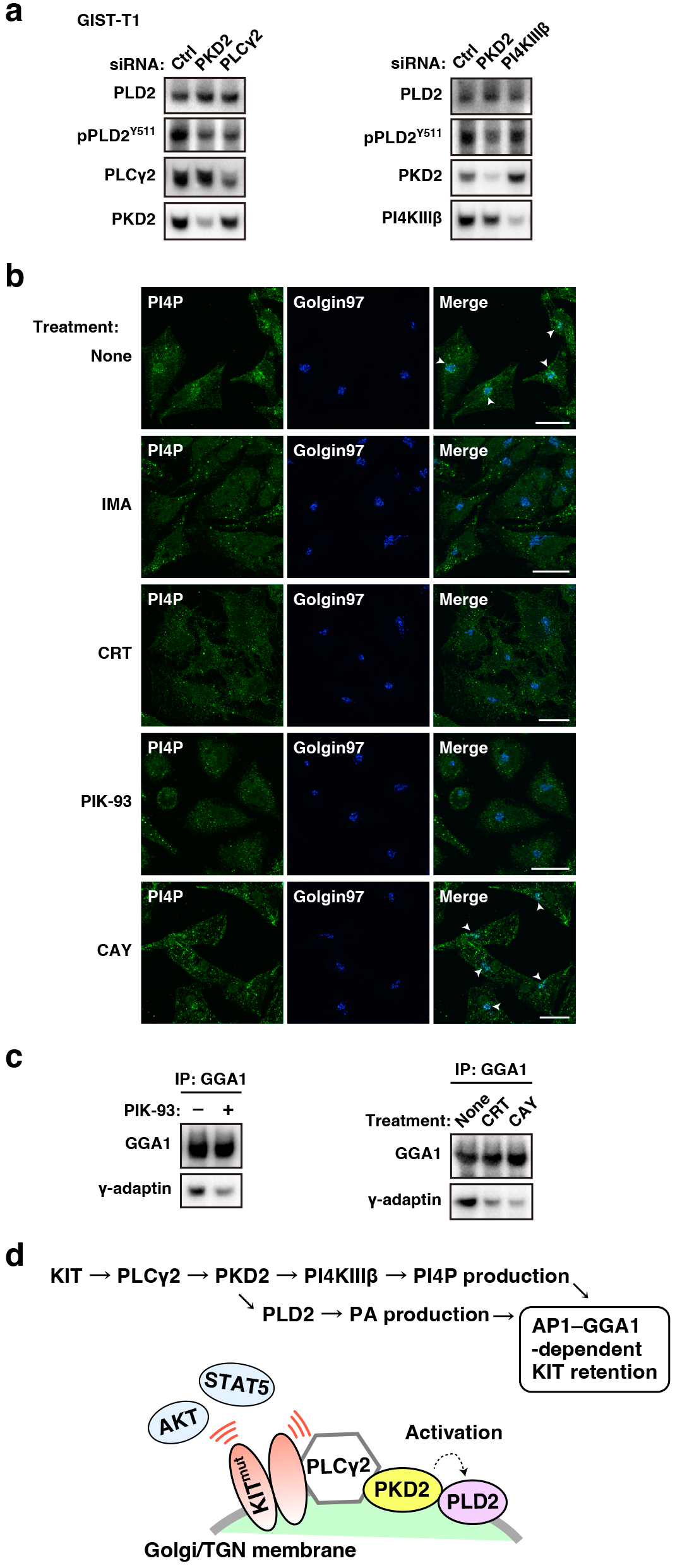
In GIST-T1 cells, PKD2, but not Pl4KIIIJ3, is an upstream molecule of PLD2. **(a)** GIST-T1 cells were transfected with the indicated siRNAs and cultured for 30 h. Lysates were immunoblotted with the indicated antibodies. pPLD2^Y511^, phospho-PLD2 Y511. **(b)** GIST-T1 cells were treated with 200 nM imatinib (IMA, a KIT tyrosine kinase inhibitor), 10 *µ*M CRT0066101 (CRT, a PKD inhibitor), 20 *µ*M PIK-93 (a PI4KllIβ inhibitor), or 20 *µ*M CAY10594 (CAY, a PLD inhibitor) for 4 h. Cells were immunostained with anti-PI4P (green) and golgin97 (TGN marker, magenta). Arrowheads indicate the Golgi/TGN area. Scale bars, 20 *µ*m. Note that PI4P in the Golgi/TGN area was decreased by treatment of imatinib, CRT0066101, and PIK-93, whereas it was retained in the presence of CAY10594. **(c)** GIST-T1 cells were treated with 20 *µ*M PIK-93, 10 *µ*M CRT0066101, or CAY10594 for 8 h. GGA1 was immunoprecipitated with anti-GGA1 antibody. The immunoprecipitates were immunoblotted for GGA1 and y-adaptin. IP, immunoprecipitation. **(d)** PKD2 separately activates PLD and PI4KllIβ in GIST cells. Both enzymatic products, PI4P and phosphatidic acid (PA), directly or indirectly play a role in the association of GGA1 with AP1, which is important for Golgi/TGN retention of KIT^mut^ If these pathways are blocked, KIT^mut^ is released from the Golgi/TGN region, resulting in degradation in lysosomes.

We found that the imatinib-resistant GIST cell lines, GIST-R9 and GIST430, expressed PLD1, unlike GIST-T1 cells. PLD2 knockdown partially decreased the protein levels of KIT in these cells compared with PLD2-knocked down GIST-T1 cells (Suppl. Fig. 3a). In GIST430 cells, the knockdown of PLD1 decreased KIT by 10%, indicating that the involvement of PLD1 in KIT retention depends on the cell context. AKT and STAT5 phosphorylation was not reduced in PLD1- or PLD2-knocked down cells. Although pSTAT5 was partially decreased, simultaneous knockdown of PLD1 and PLD2 in GIST-R9 and GIST430 cells did not affect KIT levels or pAKT to a similar extent as in GIST-T1 cells (Suppl. Fig. 3b). These imatinib-resistant GIST cells may compensate for the loss of PLD1 and PLD2 proteins and restore PA levels in PLD1/PLD2-knocked down cells. At present, we cannot explain the reason for this. Further studies are required to elucidate these compensation mechanisms.

## Discussion

In this study, we demonstrated that PLD activity was required for Golgi/TGN retention by KIT^mut^ in GIST (Fig. 4d). The PLD inhibitors CAY10594 and 1-butanol move KIT^mut^ from the Golgi/TGN, resulting in downstream inactivation. After release from the Golgi/TGN region, KIT^mut^ is incorporated into lysosomes for degradation. PLD inhibitors have a similar effect as PKD inhibitors on KIT^mut^ localisation and growth signalling in GIST cells. KIT^mut^ activates PLD2 via the PLCγ2–PKD2 pathway. PKD2 could separately activate PLD2 and PI4KIIIβ for producing PA and PI4P, respectively. Both PLD2 and PI4KIIIβ are required for the association of AP1 with GGA1, which is a cause of Golgi retention of KIT^mut^ in GIST cells. The loss-of-function of PLD does not affect the signalling of other RTK^mut^ mutations such as FLT3-ITD in AML. These findings suggest that KIT^mut^ separately activates PLD2 and PI4KIIIβ through the PLCγ2–PKD2 pathway for its Golgi retention in GIST cells.

Previous studies have demonstrated Golgi/TGN retention by RTKs other than KIT^mut^ and FLT3-ITD. Constitutively active forms of PDGFRA, FGFR3, and TRKA are found in the Golgi/TGN region and are autophosphorylated in cancer cells^40–44^. In hepatocellular carcinoma cells, overexpressed MET was retained in the Golgi/TGN region in a manner dependent on its tyrosine kinase activity^45^. Tyrosine phosphorylates the HER3 kinase, leading to the autonomous proliferation of host cells. Non-receptor-type oncogenic proteins, such as Src-family tyrosine kinases, RAS GTPases, and mammalian target of rapamycin, have been suggested to use the Golgi/TGN membrane as their signalling platform^46–51^. In particular, PLD is located downstream of RAS^52, 53^. Investigating whether RAS uses PLD for Golgi/TGN localisation is attractive. Therefore, it is of substantial interest to determine whether PLD plays a role in Golgi/TGN retention of oncogenic signal transduction proteins other than GIST KIT^mut^.

PLD1 and PLD2 generate PA and choline from PC located in the lipid bilayer^37^. PA plays multiple roles in various cellular events, including lipid biogenesis, membrane trafficking, cytoskeletal reorganisation, and acts as a scaffold for signal transduction^54–56^. In the transport carrier formation process at the Golgi/TGN membrane, PA provides platforms for membrane trafficking proteins and creates membrane curvature owing to its cone-like shape^26–28^. In GIST cells, autophosphorylation of KIT in the Golgi/TGN membrane, which does not occur in normal cells, aberrantly activates PLD2 via the PLCγ2–PKD2 pathway, probably resulting in local dysregulated PA production. We hypothesised that abnormal local PA production would stop the export of KIT^mut^ from the Golgi/TGN region. Abnormally generated PA may play a role in the induction of the AP1–GGA1 interaction, which is involved in KIT retention. Analysis of PA distribution with fluorescence imaging and quantification of PA amounts will help us understand the precise mechanism of PA-dependent Golgi/TGN retention by KIT^mut^ in GIST cells.

In the present study, we identified PLD2 as a protein downstream of KIT^mut^ in GIST. Previously, several groups reported that in mast cells, SCF-stimulated normal KIT at the PM induces PLD activation, resulting in chemical mediator release^57, 58^. These studies suggest that the SCF–KIT axis physiologically activates PLD in haematopoietic cells, melanocytes, germ cells in the testes, and interstitial Cajal cells. Therefore, there is great interest in examining the physiological role of PLD in these cells.

Previous studies reported that ADP-ribosylation factor (ARF) proteins act as mediators between PKD and PLD^17–19, 59, 60^. In our recent report, we knocked down Golgi-localised ARFs such as ARF1, ARF4, and ARF5 in GIST-T1 cells^61^. However, knockdown of ARF1, ARF4, or ARF5 did not phenocopy PLD2 knockdown. Furthermore, since the simultaneous knockdown of ARF1, ARF4, and ARF5 blocks the ER export of KIT^mut^, a conclusion cannot be drawn from a simple ARF loss-of-function study. Cell-free system methods and protein–protein interaction analyses will be powerful tools for understanding the precise mechanism of PLD activation through ARF function.

In imatinib-resistant GIST cell lines, knockdown of PLD1 and PLD2 could not fully explain the effect of the PLD inhibitor CAY10594. Genetic alterations occur not only in the *KIT* gene^62^. Multiple enzymes such as diacylglycerol kinases, lysophosphatidic acid acyltransferase, PA phosphatases, and phospholipase A are involved in controlling intracellular PA levels^28, 63^. In GIST-R9 and GIST430 cells, alterations in the expression of PA-related proteins may compensate for the loss of PLD1 and PLD2 by siRNA. Further studies are required to understand the mechanisms underlying Golgi retention by KIT^mut^ in imatinib-resistant cells.

In conclusion, we showed that in addition to PI4KIIIβ, PLD2 is an effector protein of the KIT–PLCγ2–PKD2 cascade, and its enzymatic product PA may be a key player for KIT retention in the Golgi/TGN in GIST cells. PLD2 maintains the uncontrolled growth signalling of KIT^mut^ by trapping the mutant in the Golgi/TGN. Our findings provide new insights into the role of PLD2 in GIST growth. Moreover, from a clinical perspective, our findings offer a new strategy for GIST treatment by releasing KIT^mut^ from the signalling platform.

## Materials and Methods

### Cell culture

GIST-T1 cells (Cosmo Bio, Tokyo, Japan) and GIST-R9 cells were cultured at 37°C in Dulbecco’s modified Eagle’s medium (DMEM) supplemented with 10% fetal calf serum (FCS), penicillin, and streptomycin (Pen/Strep). GIST-R9 cells were incubated with 1 µM imatinib as described previously^62^. GIST430/654 (hereafter referred to as GIST430) cells were kindly provided by Dr. Jonathan Fletcher (Dana-Farber Cancer Institute, Boston, MA, USA). GIST430 cells were cultured at 37°C in Iscove’s modified Dulbecco’s medium (IMDM) supplemented with 15% FCS and Pen/Strep. GIST430 cells were maintained in the presence of 100 nM imatinib^64^. Kasumi-1 (JCRB Cell Bank, Osaka, Japan), MOLM-14 (DSMZ, Braunschweig, Germany), and HMC-1.2 cells^65^ were cultured at 37°C in RPMI1640 supplemented with 10% FCS and Pen/Strep. All human cell lines were authenticated by short tandem repeat analysis at the JCRB Cell Bank and tested for *mycoplasma* contamination using a MycoAlert Mycoplasma Detection Kit (Lonza, Basel, Switzerland).

### Antibodies

The antibodies used for immunoblotting, immunoprecipitation, and immunofluorescence are listed in Supplementary Table 1.

### Chemicals

CAY10594, imatinib mesylate (Cayman Chemical, Ann Arbor, MI, USA), PIK-93, and CRT0066101 (Selleck Chemicals) were dissolved in dimethyl sulfoxide. NH_4_Cl and 1-butanol were purchased from Sigma-Aldrich (St. Louis, MO) and Fujifilm Wako Chemicals (Osaka, Japan), respectively.

### Gene silencing with siRNA

For silencing, *PLD1*, *PLD2*, *PLCγ2*, *PKD2*, and *PI4KIIIβ* ON-TARGETplus SMARTpool siRNAs were purchased from Horizon Discovery (Waterbeach, UK). A list of the siRNAs used in this study is provided in Supplementary Table 2. Electroporation was performed using the NEON Transfection System (Thermo Fisher Scientific) according to the manufacturer’s instructions.

### Immunofluorescence confocal microscopy

GIST-T1 cells were cultured on poly L-lysine-coated coverslips and fixed with 4% paraformaldehyde (PFA) for 20 min at room temperature. The fixed cells were permeabilised and blocked for 30 min in Dulbecco’s phosphate-buffered saline (D-PBS(−)) supplemented with 0.1% saponin and 3% bovine serum albumin (BSA), and then incubated with primary and secondary antibodies for 1 h each. After washing with D-PBS(−), cells were mounted using Fluoromount (Diagnostic BioSystems, Pleasanton, CA, USA). Confocal images were obtained using a FLUOVIEW FV3000 (Olympus, Tokyo, Japan) or TCS SP8 (Leica, Wetzlar, Germany) laser scanning microscope. Composite figures were prepared using FV31S-SW (Olympus), Leica Application Suite X Software (Leica), Photoshop, and Illustrator (Adobe, San Jose, CA, USA).

### Immunostaining for PI4P visualisation

GIST-T1 cells were treated with 50 mM NH_4_Cl in D-PBS(−) before fixation with 2% PFA. Permeabilisation and staining were performed as previously described. To stain for PI4P, anti-PI4P mouse IgM (Echelon Biosciences, Salt Lake City, UT, USA) and goat Alexa Fluor 488-conjugated anti-mouse IgM (Thermo Fisher Scientific) were used as primary and secondary antibodies, respectively. After washing with D-PBS(−), stained cells were fixed with 2% PFA for 15 min, treated with 50 mM NH_4_Cl in D-PBS(−), and washed with double distilled water.

### Western blotting

Lysates prepared in sodium dodecyl sulphate-polyacrylamide gel electrophoresis (SDS-PAGE) sample buffer were subjected to SDS-PAGE and electrotransferred onto polyvinylidene fluoride membranes. Briefly, 5% skim milk in Tris-buffered saline containing Tween 20 (TBST) was used to dilute the antibodies. For immunoblotting with anti-pKIT Y703, anti-pPLD2 Y169 or Y511, the antibodies were diluted with 3% BSA in TBST. Immunodetection was performed using the Immobilon Western Chemiluminescent HRP Substrate (Sigma-Aldrich). Sequential re-probing of the membranes was performed after complete removal of antibodies with Restore PLUS Western Blot Stripping Buffer (Thermo Fisher Scientific) and/or inactivation of peroxidase by 0.1% sodium azide. The results were analysed using a ChemiDoc XRC+ with Image Lab software (Bio-Rad, Hercules, CA, USA).

### Immunoprecipitation

Lysates from imatinib-treated GIST-T1 cells were prepared in NP-40 lysis buffer supplemented with 50 mM HEPES (pH 7.4), 10% glycerol, 1% NP-40, 4 mM EDTA, 100 mM NaF, 1 mM Na_3_VO_4_, protease inhibitor cocktail, 2 mM β-glycerophosphate, 2 mM sodium pyrophosphate, and 1 mM phenylmethylsulfonyl fluoride. Immunoprecipitation was performed at 4°C for 5 h using Dynabeads Protein G beads pre-coated with anti-KIT, anti-GGA1, or anti-PLD2 antibodies. Immunoprecipitates were dissolved in SDS-PAGE sample buffer and immunoblotted.

### PLD2-MYC transfection

A gene encoding carboxy-terminal MYC-tagged PLD2 (PLD2-MYC), inserted into the pRP vector for expression in mammalian cells, was purchased from VectorBuilder (Yokohama, Japan; vector ID, VB230228-1368ssm). Transient transfection of PLD2-MYC was performed using PEIMAX (Polysciences, Warrington, PA, USA) or JetOPTIMUS (Polyplus-Transfection, Illkirch, France). PLD2-MYC expression was detected by immunoblotting and immunofluorescence using an anti-MYC antibody (71D10).

## Acknowledgments

This work was supported by a grant-in-aid for Scientific Research from the Japan Society for the Promotion of Science, Japan (24K10370 to Y. O. and 22K08883 to T. N.), a research grant from the Research Foundation for Pharmaceutical Sciences, Japan (to Y. O.), the Japan Foundation for Applied Enzymology, Japan (to Y. O.), the Life Science Foundation of Japan (to Y. O.), and the Foundation for the Promotion of Cancer Research, Japan (to Y. O.).

## Author contributions

Y. O. conceived, designed, performed, and analysed the data from all experiments and wrote the manuscript. M. N. performed the immunoblotting and edited the manuscript. I. S. conceived and supervised the project and analysed the data. T. T. provided advice regarding the design of the *in vitro* experiments. T. N. provided advice on the design of the *in vitro* experiments and edited the manuscript. All the authors have read and approved the final version of the manuscript.

## Funding

The Japan Society for the Promotion of Science (24K10370 to Y. O. and 22K08883 to T. N.). Research Foundation for Pharmaceutical Sciences (to Y. O.). Japan Foundation for Applied Enzymology (Y. O.). Life Science Foundation of Japan (to Y. O.). Foundation for the Promotion of Cancer Research (to Y. O.).

## Competing interests

The authors declare no competing interests.

## Abbreviations

AML: acute myelogenous leukemia
AP1: adaptor protein-1 complex
ARF: ADP-ribosylation factor
BSA: bovine serum albumin
CAY: CAY10594
CRT: CRT0066101
DMEM: Dulbecco’s modified Eagle’s medium
D-PBS(−): Dulbecco’s phosphate-buffered saline
siRNA: small interfering RNA
ER: endoplasmic reticulum
ERK: extracellular signal-regulated kinase
FCS: fetal calf serum
FGFR: fibroblast growth factor receptor
FLT3-ITD: FMS-like tyrosine kinase 3-intenal tandem duplication
GGA: Golgi-associated, γ-adaptin ear-containing, ARF-binding protein
GIST: gastrointestinal stromal tumor
IP: immunoprecipitation
LAMP1: lysosome-associated membrane protein 1
MCL: mast cell leukemia
mut: mutant
NH_4_Cl: ammonium chloride
PA: phosphatidic acid
PC-PLD: phosphatidylcholine-selective phospholipase D
PDGFR: platelet-derived growth factor receptor
Pen/Strep: penicillin and streptomycin
PFA: paraformaldehyde
PI3K: phosphatidylinositol-3 kinase
PI4KIIIβ: phosphatidylinositol-4 kinase IIIβ
PI4P: phosphatidylinositol 4-phosphate
PKD: protein kinase D
pKIT: phospho-KIT
PLCγ: phospholipase Cγ
PM: plasma membrane
RTK: receptor tyrosine kinase
SCF-R: stem cell factor receptor
SDS-PAGE: sodium dodecyl-sulfate polyacrylamide gel electrophoresis
siRNA: small interfering RNA
STAT: signal transducer and activator of transcription
TBST: Tris-buffered saline with Tween 20
TfR: transferrin receptor
TGN: *trans-*Golgi network
TKI: tyrosine kinase inhibitor
TRKA: tropomyosin receptor kinase A.

## Notes

**Grant Support:** Japan Society for the Promotion of Science (24K10370 to Y. O. and 22K08883 to T. N.) The Research Foundation for Pharmaceutical Sciences (to Y. O.) Japan Foundation for Applied Enzymology (to Y. O.) Life Science Foundation of JAPAN (to Y. O.) Foundation for Promotion of Cancer Research (to Y. O.)

### Competing Interest Statement

The authors have declared no competing interest.

### Summary of Updates

Legends for Figures 1-4 were revised.

