## Supplementary Figures 1-3 for "Golgi retention of KIT in gastrointestinal stromal tumour cells is phospholipase D activity-dependent"

### Supplementary Information

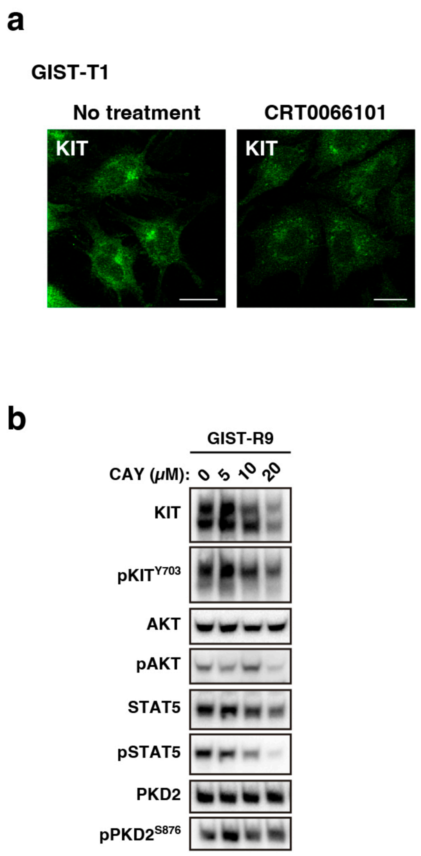


**Supplementary Figure 1. KIT$^{mut}$ is retained in the Golgi/TGN region in a PKD2-dependent manner in GIST-T1 cells.**
(**a**) GIST-T1 cells were treated with 20 $\mu$M CRT0066101 (a PKD inhibitor) for 4 h and then immunostained with anti-KIT antibody. Scale bars, 20 $\mu$m. (**b**) GIST-R9 cells were treated with CAY10594 (CAY, a PLD inhibitor) for 8 h and then immunoblotted. pKIT$^{Y703}$, phospho-KIT Y703; pPKD2$^{S876}$, phospho-PKD2 S876.

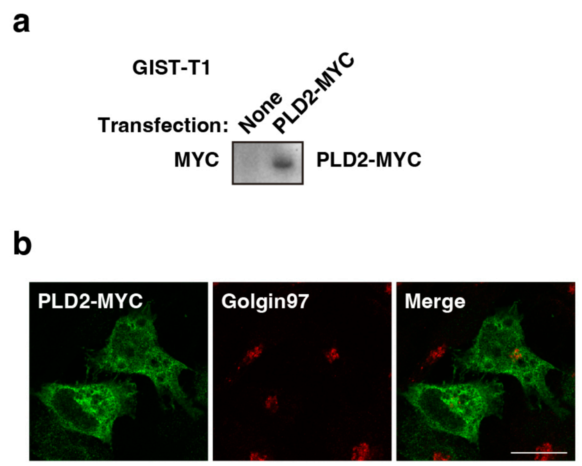


**Supplementary Figure 2. Localisation of PLD2 in GIST-T1 cells.**

(**a**,**b**) GIST-T1 cells were transfected with PLD2-MYC for 24 h. (**a**) Lysates were immunoblotted with an anti-MYC antibody. (**b**) Cells were immunostained with anti-MYC (green) and anti-golgin97 (red) antibodies. Scale bar, 20 $\mu$m.

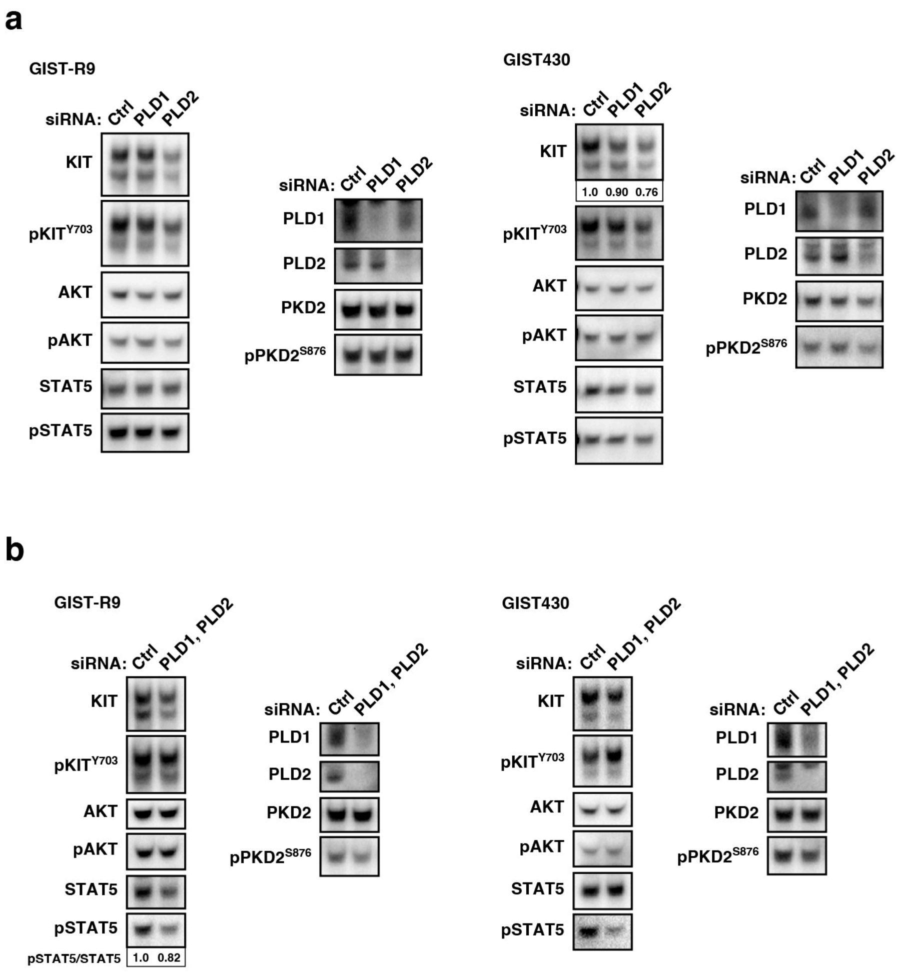


**Supplementary Figure 3. Effect of knockdown of PLD in GIST-R9 cells and GIST430 cells.**

(**a**,**b**) GIST-R9 cells and GIST430 cells were transfected with the indicated siRNAs for 48 h. Lysates were immunoblotted with the indicated antibodies. pPKD2$^{S876}$, phospho-PKD2 S876. (**a**) Relative band intensities normalized with each control sample are shown. (**b**) Levels of pSTAT5 are expressed relative to control cell sample, after normalization with STAT5.

**Supplementary Table 1. List of antibodies**

| Antibody | Distribution source | Identifier |
|---|---|---|
| AKT (40D4) Mouse mAb | Cell Signaling Technology | Cat#2920; PRID: AB_1147620 |
| AKT (clone 55) Mouse mAb | BD Transduction Laboratories | Cat#610861 RRID:AB_398180 |
| Phospho-AKT (T308) (C31E5E) Rabbit mAb | Cell Signaling Technology | Cat#2965; PRID: AB_2255933 |
| β-actin Rabbit polyclonal ab | Cell Signaling Technology | Cat#4967; PRID: AB_330288 |
| FLT3 (8F2) Rabbit mAb | Cell Signaling Technology | Cat#4967; PRID: AB_330288 |
| γ-Adaptin (clone 88) Mouse mAb | BD Transduction Laboratories | Cat#610386; PRID: AB_397768 |
| GGA1 Rabbit polyclonal ab | Proteintech | Cat#25674-1-AP; PRID: AB_2880188 |
| Golgin-97 (D8P2K) Rabbit mAb | Cell Signaling Technology | Cat#13192; PRID: AB_2798144 |
| Golgin-97 (CDF4) Mouse mAb | Thermo Fisher Scientific | Cat#14-9767-82; PRID: AB_2573010 |
| KIT (D13A2) Rabbit mAb | Cell Signaling Technology | Cat#3074; PRID: AB_1147633 |
| KIT (D3W6Y) Rabbit mAb | Cell Signaling Technology | Cat#37805; PRID: AB_2799120 |
| KIT (E-1) Mouse mAb | Santa Cruz Biotechnology | Cat#sc-17806; PRID: AB_626875 |
| Phospho-KIT (Y703) (D12E12) Rabbit mAb | Cell Signaling Technology | Cat#3073; PRID: AB_1147635 |
| LAMP1 (D4O1S) Mouse mAb | Cell Signaling Technology | Cat#15665; PRID: AB_2798750 |
| MYC-Tag (71D10) Rabbit mAb | Cell Signaling Technology | Cat#2278; PRID: AB_10828091 |
| PI4KIIIβ (clone 7) Mouse mAb | BD Transduction Laboratories | Cat#sc-17806; PRID: AB_626875 |
| PI4P Mouse mAb | Echelon Biosciences | Cat#Z-P004; PRID: AB_11127796 |
| PKD2 (D1A7) Rabbit mAb | Cell Signaling Technology | Cat#8188; PRID: AB_10829368 |
| PKD2 (O95G1) Mouse mAb | BioLegend | Cat#617302; PRID: AB_2810668 |
| Phospho-PKD2 (S876) (EP1496Y) Rabbit mAb | Abcam | Cat#ab51251; PRID: AB_882060 |
| PLCγ2 (E5U4T) Rabbit mAb | Cell Signaling Technology | Cat#55512; PRID: AB_2799488 |
| PLD1 Goat polyclonal ab | R&D SYSTEMS | Cat#AF5615; RRID:AB_2163858 |
| PLD1 (F-12) Mouse mAb | Santa Cruz Biotechnology | Cat#sc-28314; RRID:AB_677324 |
| PLD1 Rabbit mAb | Abcam | Cat#ab68150 |
| PLD2 Goat polyclonal ab | R&D SYSTEMS | Cat#AF10123 |
| Phospho-PLD2 (Y169) Rabbit polyclonal ab | St John's Laboratory | Cat#STJ90989 |
| Phospho-PLD2 (Y511) Rabbit polyclonal ab | Thermo Fisher Scientific | Cat#PA5-105377; PRID: AB_2816805 |
| STAT5 (C-17) Rabbit polyclonal ab | Santa Cruz Biotechnology | Cat#sc-835; PRID: AB_632446 |
| STAT5 (clone 89) Mouse mAb | BD Transduction Laboratories | Cat#610192; PRID: AB_397590 |
| STAT5 (D2O6Y) Rabbit mAb | Cell Signaling Technology | Cat#94205; PRID: AB_2737403 |
| STAT5 [pY694] Rabbit mAb | Cell Signaling Technology | Cat#4322; PRID: AB_10544692 |
| Transferrin Receptor (H68.4) Mouse mAb | Thermo Fisher Scientific | Cat#13-6800; PRID: AB_2533029 |
| HRP donkey anti-mouse IgG | Jackson ImmunoResearch | Cat#715-035-151; PRID: AB_2340771 |
| HRP donkey anti-rabbit IgG | Jackson ImmunoResearch | Cat#711-035-152; PRID: AB_10015282 |
| HRP donkey anti-goat IgG | Jackson ImmunoResearch | Cat#705-035-147; PRID: AB_2313587 |
| Donkey anti-Rabbit IgG (H+L), Alexa Fluor 488 | Thermo Fisher Scientific | Cat#A-21206; PRID: AB_2535792 |
| Donkey anti-Mouse IgG (H+L), Alexa Fluor 568 | Thermo Fisher Scientific | Cat#A-10037; PRID: AB_2534013 |
| Donkey anti-Rabbit IgG (H+L), Alexa Fluor 647 | Thermo Fisher Scientific | Cat#A-31573; PRID: AB_2536183 |
| Goat anti-Mouse IgM (Heavy chain), Alexa Fluor 488 | Thermo Fisher Scientific | Cat#A-21042; PRID: AB_2535711 |

#### Supplementary Table 2. List of siRNAs

| Oligonucleotides | Distribution source | Cat# |
|---|---|---|
| **ON-TARGETplus Human PKD2 (25865) siRNA - SMARTpool** | **Horizon Discovery** | **L-004197-00** |
| **ON-TARGETplus Human PLCγ2 (5336) siRNA - SMARTpool** | **Horizon Discovery** | **L-008339-02** |
| **ON-TARGETplus Human PI4KIIIβ (5298) siRNA - SMARTpool** | **Horizon Discovery** | **L-006777-00** |
| **ON-TARGETplus Human PLD1 (5337) siRNA – SMARTpool** | **Horizon Discovery** | **L-009413-00** |
| **ON-TARGETplus Human PLD2 (5338) siRNA – SMARTpool** | **Horizon Discovery** | **L-005064-00** |
| **ON-TARGETplus Non-targeting Pool** | **Horizon Discovery** | **D-001810-10-20** |
